# Probing different conformational states of polar organizing protein Z with AlphaFold

**DOI:** 10.1101/2023.10.11.561952

**Authors:** Ishan Taneja, Keren Lasker

**Affiliations:** Department of Integrative Structural and Computational Biology, Scripps Research, La Jolla, CA 92037

## Abstract

AlphaFold has been a remarkably powerful tool to predict the structures of proteins. Here, we demonstrate that AlphaFold is capable of sampling different conformational states of the oligomerization domain of polar organizing protein Z (PopZ). More broadly, our work suggests that conformational heterogeneity can be extracted from AlphaFold without directly modifying the input multiple sequence alignment.

## Introduction

AlphaFold (AF) has demonstrated remarkable success in its ability to predict the three-dimensional structures of proteins with sufficiently deep multiple sequence alignments (MSAs). However, whether conformational heterogeneity can be recovered from AlphaFold remains an active area of research. Current methods that predict different conformational states of a protein generally do so by manipulating the input MSA^1–3^. For example, Steele et. al clustered the input MSA and then accordingly subsampled it to recover different folds of metamorphic proteins^1^. Likewise, del Alamo et. al restricted the depth of randomly subsampled the MSA to predict diverse conformational ensembles of transporters and receptor proteins^2^. In a different context (but similar methodological theme), Bryant et. al modified the AF-Multimer pipeline by adding a learnable bias parameter to the input MSA per protein complex to improve its model confidence score^4^.

While these methods have shown promising results, we developed a complementary method to probe different conformational states. Our preliminary approach leverages a parameter-efficient fine-tuning method to learn a low-dimensional parameter that can be added to the model weights to transform an initial protein structure prediction *s*_1_ to a different conformation *s*_2_. We then stochastically perturb this low-dimensional parameter by performing a random walk to generate new conformational states. Our method generates diverse conformations of the oligomerization domain of polar organizing protein Z (PopZ)^5^ and are in line with motions observed by all-atom molecular dynamics (MD) simulations.

## Methods

An objective function’s intrinsic dimension represents the lowest dimensional subspace in which one can optimize the original objective function to within a certain level of approximation error ^6,7^. This idea has been used to analyze and fine-tune large-language models (LLMs) ^7,8^. The idea is to perturb the parameters of each layer of the model via 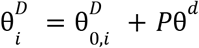, where 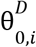 are the initial parameters of layer *i, p* is a random projection matrix defined by the Fastfood transform ^9^ mapping ℝ^*D*^ to ℝ^*d*^, and θ^*d*^ is the learned low-dimensional parameter (where *d* < *D*). We utilized this idea to derive θ^*d*^ by training OpenFold’s PyTorch implementation of AF^10^ to predict a single different conformation of PopZ derived from a MD simulation. We restricted the layers to only those in the evoformer module, with *d* = 10^4^. The model was trained with the Adam optimizer^11^, a learning rate of 1e-3, and for 200 consecutive epochs with the standard AF loss function. For reference, 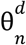 refers to the low-dimensional parameter learned at epoch *n* and 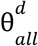 refers to set of low-dimensional parameters learned across all epochs (i.e 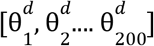).

To generate a candidate conformation, we performed a random walk in a *d*-dimensional subspace, where 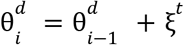, where 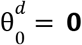 and ξ^*t*^ ∼ *N*(**0**, Σ). Σ was set to a random covariance matrix via the transformation *diag*(*S*) * *R* * *diag*(*S*), where *R* is a random correlation matrix generated according to the method of Pourahamdi et. al^12^, and *S* is vector of standard deviations drawn from 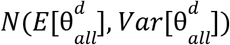, with the exception that if a diagonal entry was less than 0, it was set to 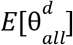. Because sampling from high-dimensional multivariate normal distributions is computationally expensive, we generated samples via *Lx* where *L* is a lower-triangular matrix generated by the Cholesky decomposition of Σ and *x* is a *d*-dimensional vector, where each entry is drawn from a standard normal distribution. The proposed conformation was accepted with probability 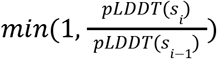, where pLDDT corresponds to the mean pLDDT score of each conformation, unless the value was less than 70, in which case it was set to 1e-10. Finally, if we observed more than three rejected steps in a row, we restarted the sampling procedure from 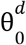. In total, this process was run for 1000 total steps.

MD simulations were run using GROMACS with the CHARMM36m force field with the corresponding CHARMM36 TIP3P water model ^13,14^. The simulations were performed at 300 K and 1 bar in the NPT ensemble using the Bussi-Donadio-Parrinello thermostat and Parrinello-Rahman barostat with a timestep of 2.0 fs. Bond lengths were constrained using LINCS. Non-bonded interactions were cut off at 1.2 Å, and long-range electrostatic interactions were calculated using the particle-mesh Ewald method with B-spline interpolation of order 4. The initial structure was sourced from an AF prediction of the sequence VAEQLVGVSAASAAASA FGSLSSALLMPKDGRTLEDVVRELLRPLLKEWLDQNLPRIVETKVEEEVQRISRGRGA. Two independent simulations were run, each for approximately .5 microseconds.

## Results

From the MD simulations, we extracted two conformations that exhibited significant variation from the initial prediction of the oligomerization domain of PopZ. The oligomerization domain of PopZ is composed of three-helix bundles, where the helix furthest away from the C-terminus (H1) is connected to the remaining two helices (H2, H3) by a flexible loop (Fig. 1). In the MD simulations, H2-H3 forms a flexible hinge enabling H3 to rotate clockwise towards H2, while H1 rotates clockwise around H2-H3 (Fig. 1). 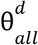 was generated by training the model to predict the magenta conformation (MD conformation 1), which was then used to sample different conformations according to the procedure described in Methods.

**Figure 1.**
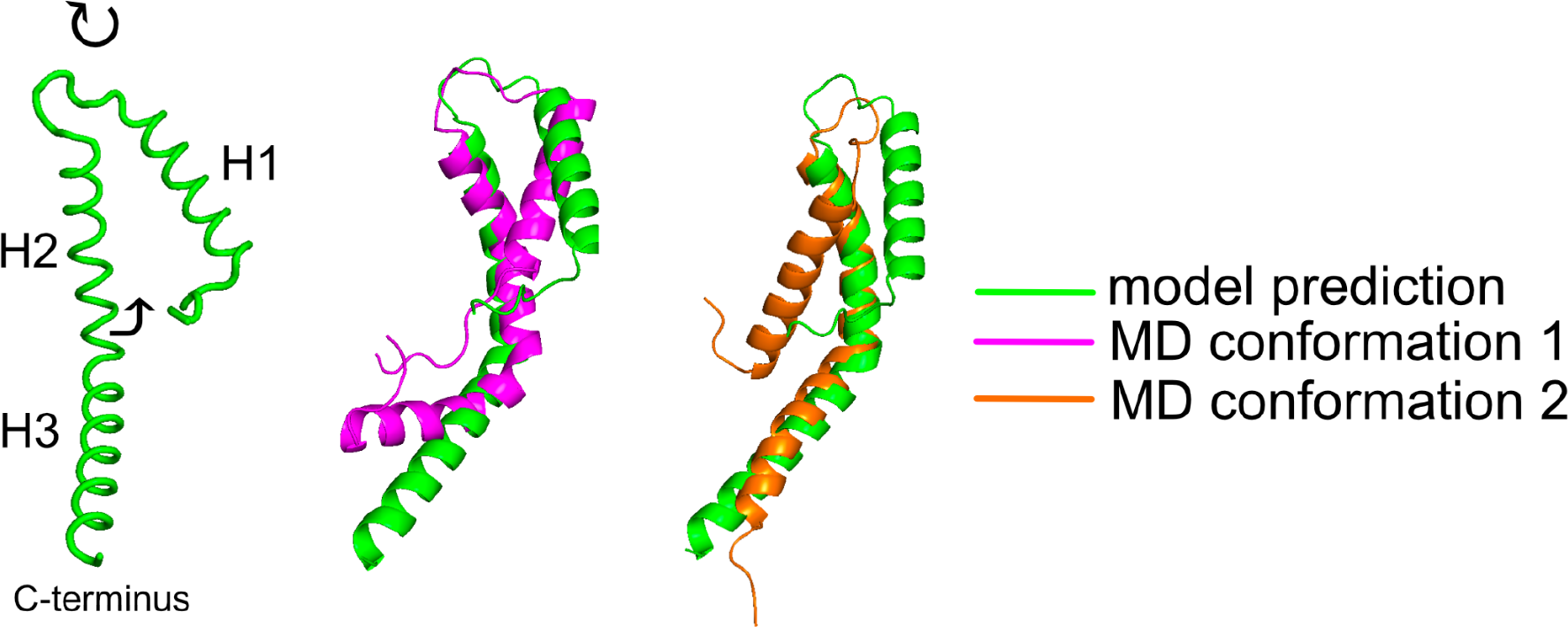
Overview of PopZ oligomerization domain conformations. Green corresponds to the initial structure predicted by the model, while the magenta conformation and the orange conformation are derived from the MD simulations.

We summarize the results of the sampling process in Figs. 2-4. Among all candidate conformations that were accepted (680 total), we projected their cartesian coordinates onto their first two principal components. Each point was either colored by its percentage helicity (Fig. 2), pLDDT score (Fig. 3), or RMSD with respect to the initial prediction (Fig. 4). In Fig. 2, we label a subset of points with their corresponding structures. Overall, we observe a heterogenous set of conformations, exhibiting variability in terms of H3’s orientation with respect to H2 and H1’s orientation with respect to H2-H3. In addition, we observe significant diversity in terms of the conformations’ secondary structure profiles, with the H1-loop region ranging from being either completely helical, partially helical, or disordered.

**Figure 2.**
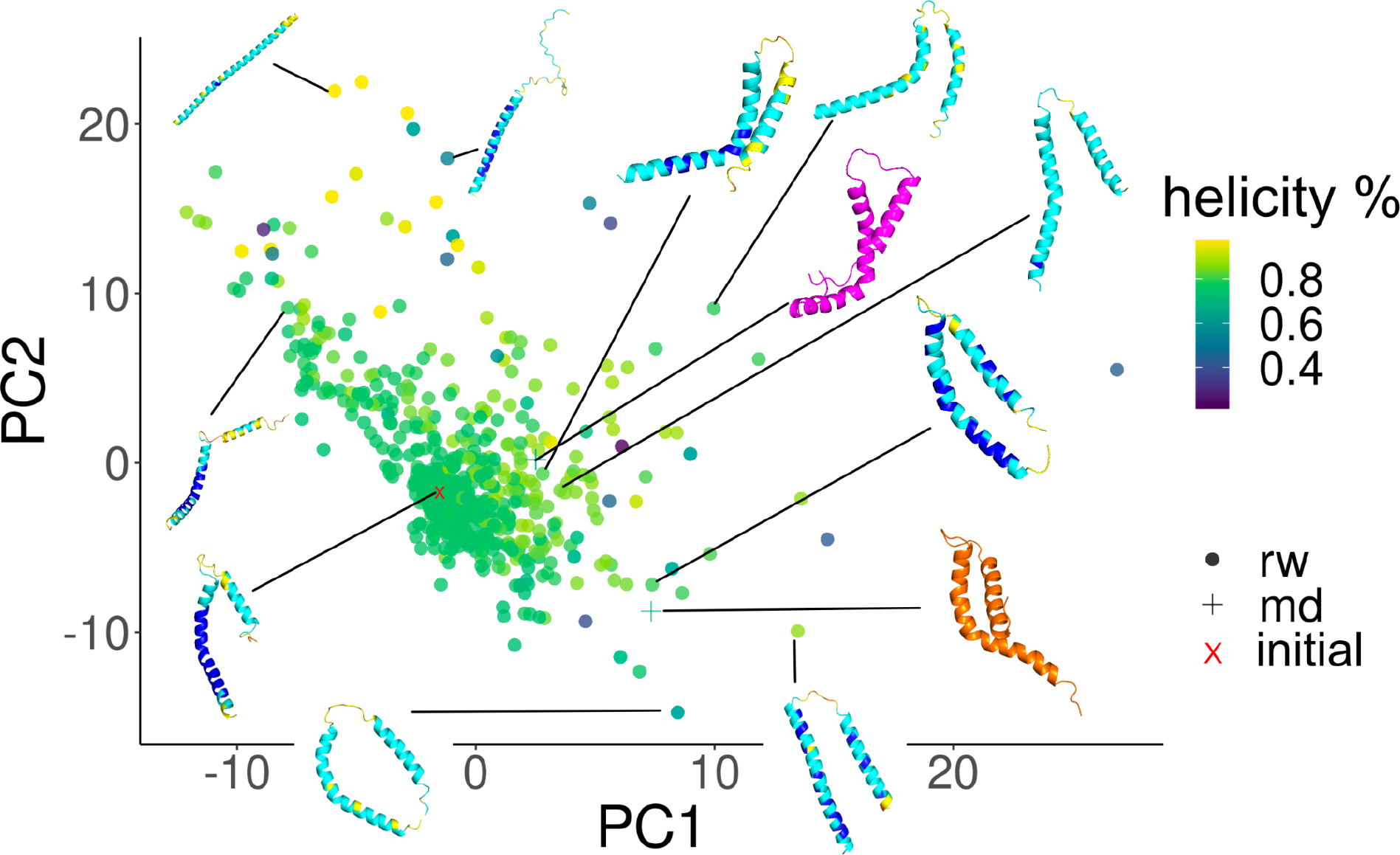
Sampled conformations projected onto their first two principal components. Each point is colored by its percentage helicity. Circular points correspond to conformations sampled by our random walk procedure, x corresponds to the initial structure predicted by the model, and + corresponds to conformations derived from the MD simulations.

**Figure 3.**
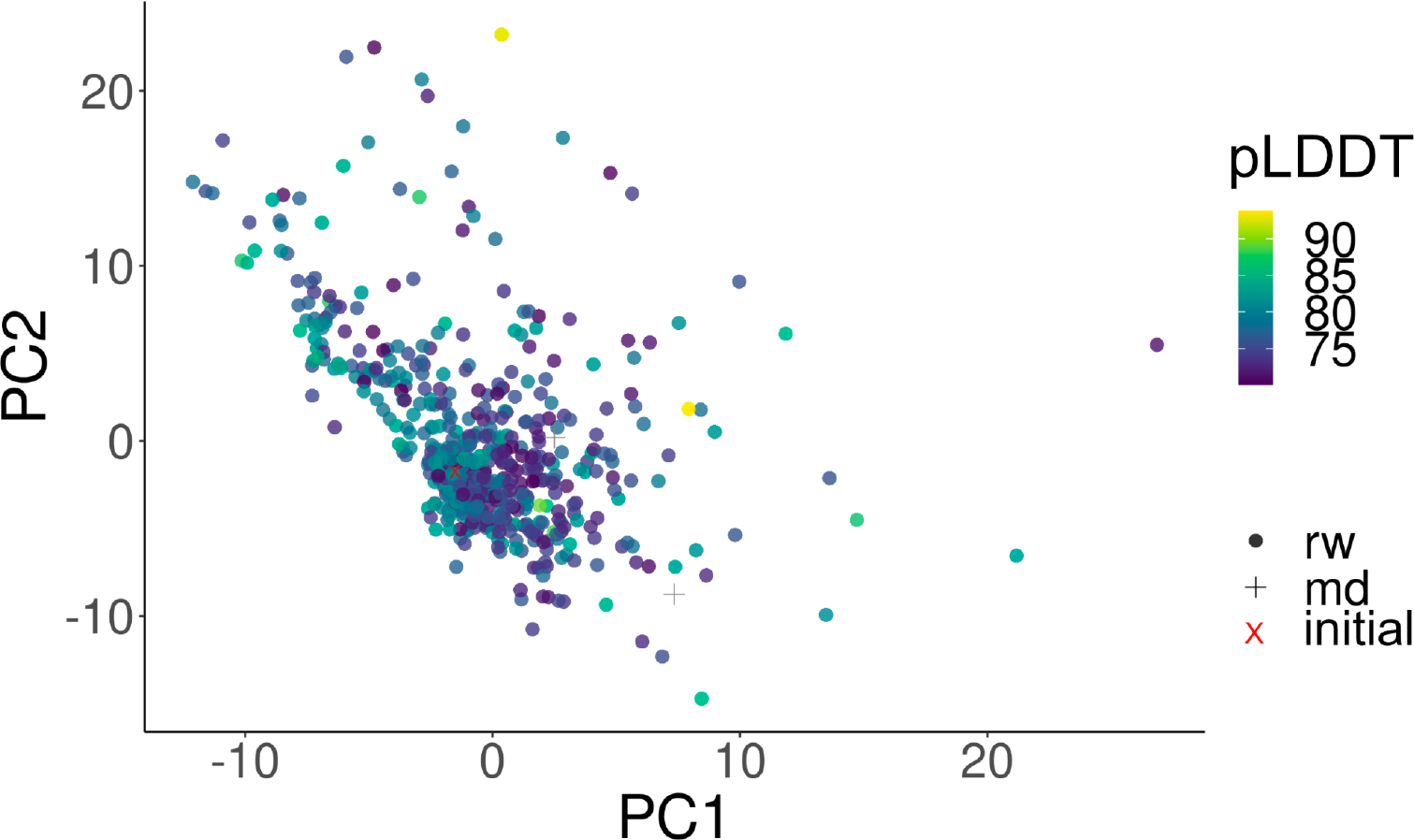
Sampled conformations projected onto their first two principal components. Each point is colored by its pLDDT score. Circular points correspond to conformations sampled by our random walk procedure, x corresponds to the initial structure predicted by the model, and + corresponds to conformations derived from the MD simulations.

**Figure 4.**
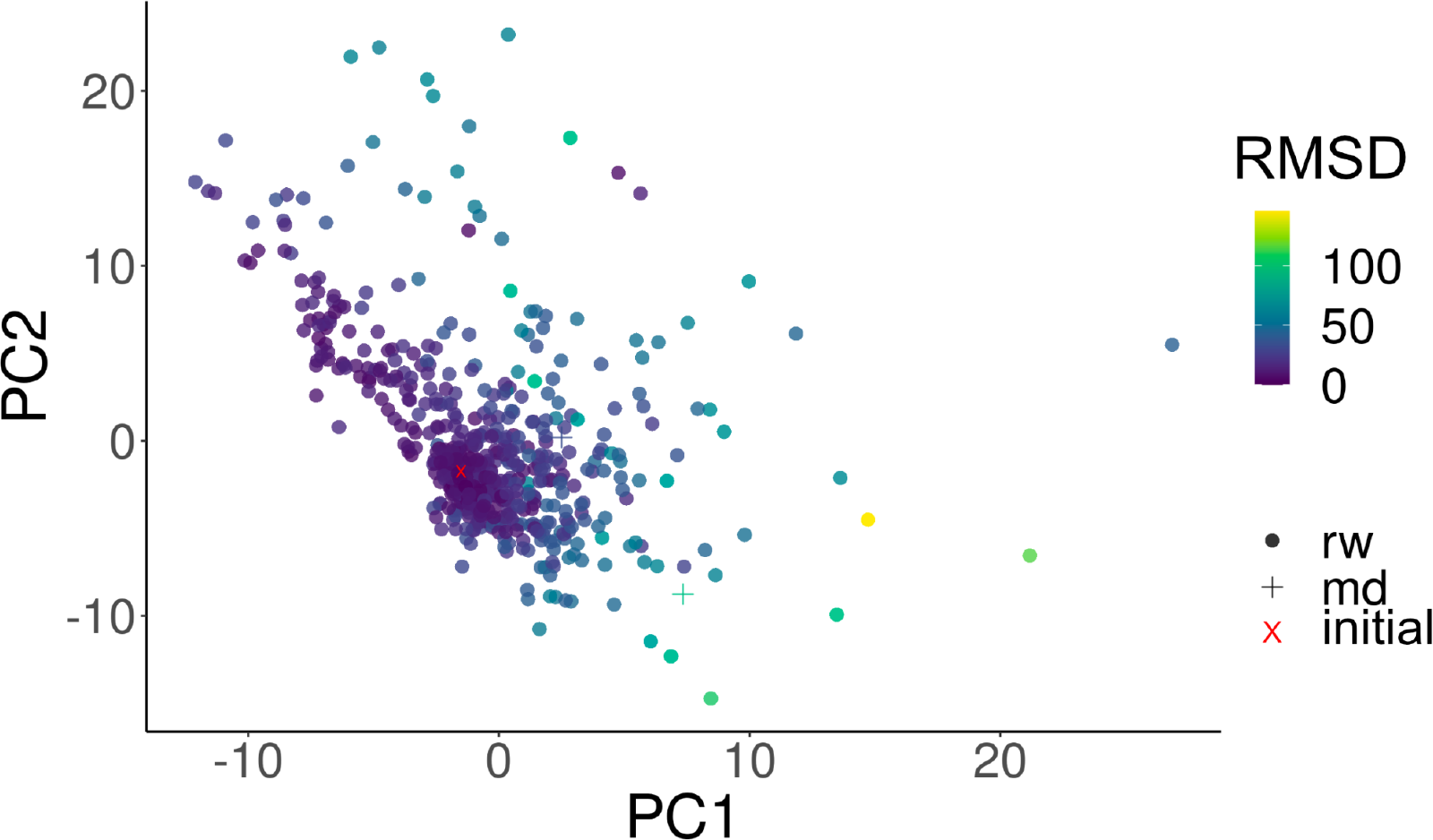
Sampled conformations projected onto their first two principal components. Each point is colored by its RMSD with respect to the initial prediction (i.e x). Circular points correspond to conformations sampled by our random walk procedure, x corresponds to the initial structure predicted by the model, and + corresponds to conformations derived from the MD simulations.

## Acknowledgements

We thank J.C. Ducom at Scripps Research High Performance Computing for computational support.

